# Serial representation of items during working memory maintenance at letter-selective cortical sites

**DOI:** 10.1101/171660

**Authors:** Ali Bahramisharif, Ole Jensen, Joshua Jacobs, John Lisman

## Abstract

We used intracranial recordings to study brain oscillations during a working memory task. To analyze sites involved in working memory, we focused on sites at which the elevation of the broadband gamma signal depended on which letter was presented. We tested a previously proposed model according to which different items are active at different phases of the theta cycle (in different gamma cycles within the theta cycle). Consistent with this model, the theta phase of letter-induced gamma elevation during maintenance reflected the order of letter presentation. These results suggest that working memory is organized by a theta-gamma code and provide strong support for the serial representation of items held in working memory.

## Introduction

Working memory (WM) is thought to be mediated by a neural system different from that which encodes long-term memory (LTM). Functionally, the WM system has a much smaller capacity than LTM system (Cowan, 2008). Physiological experiments suggest that the two memory systems are mechanistically different: whereas LTM is stored by changes in synaptic weights, WM representations involve ongoing neuronal activity (Christophel, Hebart, & Haynes, 2012; Funahashi, Bruce, & Goldman-Rakic, 1989; Fuster & Alexander, 1971; Harrison & Tong, 2009; Lee, Simpson, Logothetis, & Rainer, 2005; Riggall & Postle, 2012). However, much remains unclear about the mechanism of working memory; notably, the network processes that allow multiple items to be actively stored and read out remain unknown. Some insight into these processes has been gained from behavioral experiments using the Sternberg paradigm (Fig. 1a). It was found that, when subjects answer whether a probe item is on a just presented list, response time (RT) varies linearly with list length. This led to the suggestion that memories are actively represented serially in WM and that these are sequentially scanned during recall (S Sternberg, 1966).

**Figure 1:**
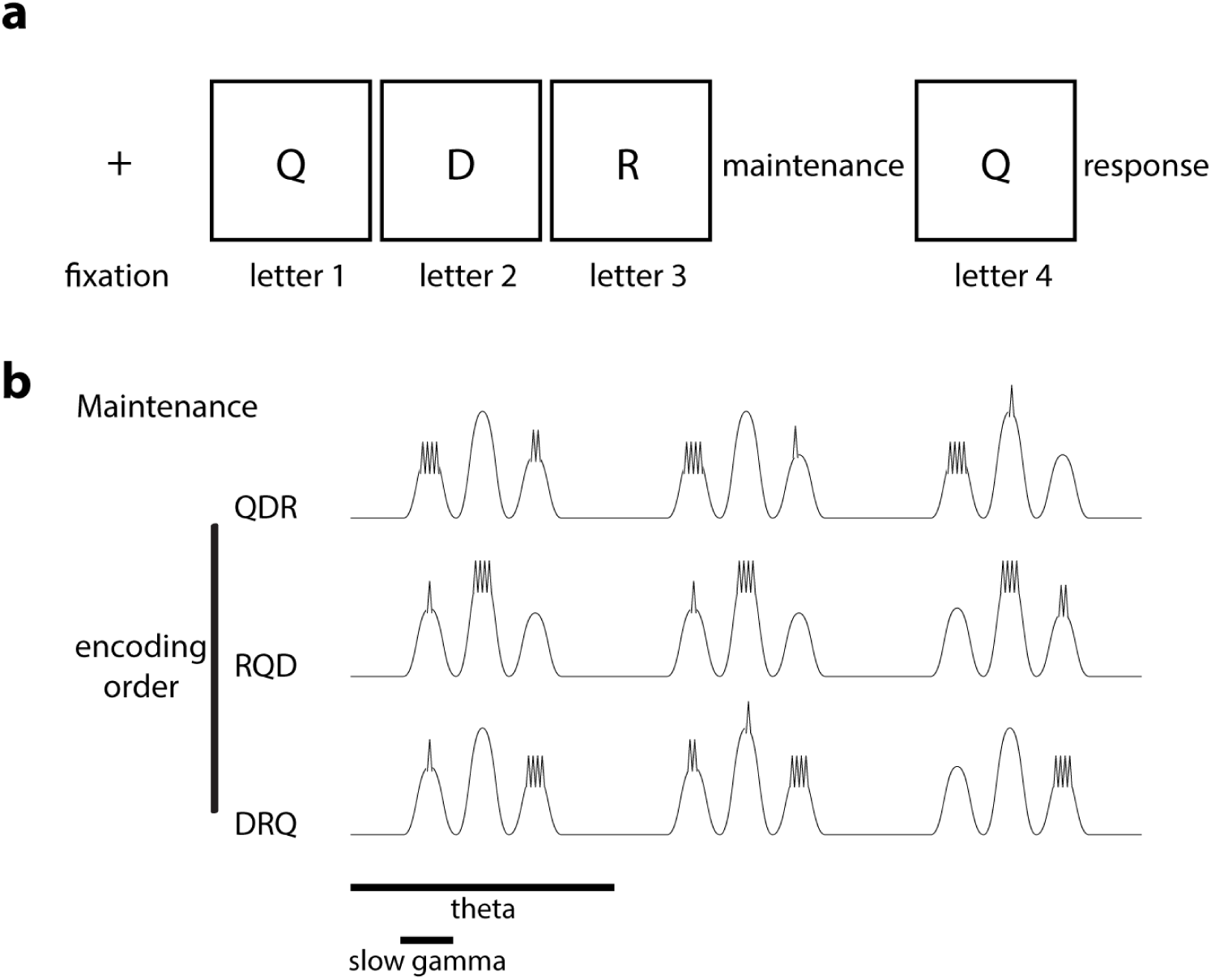
Schematic that illustrates the Sternberg task and an oscillatory model of multi-item WM. **(a)** Sternberg task with three letters each presented for 700 ms at an inter-stimulus interval of 275-350 ms. This was followed by a maintenance period of 2 s, followed by a 700 ms probe. **(b)** The LIJ model for the WM maintenance on a site selective to ‘Q’ for three encoding orders. Each theta cycle may contain many (5-8) gamma cycles (slow gamma), but only three are shown. Responses at the letter-selective sites are shown. The gamma cycle (low gamma) at which maximal high-frequency firing occurs (and thus theta phase) corresponds to the order of item presentation.

A great deal of physiological work in humans and animals has shown that the brain has oscillatory properties at multiple frequencies and that these oscillations can interact, a process called cross-frequency coupling (CFC) (Buzsáki & Wang, 2012). According to one model (Lisman/Idiart/Jensen; LIJ model) (Lisman & Jensen, 2013; Lisman & Idiart, 1995), interacting oscillations in the gamma range (>30 Hz) (reviewed in (Roux & Uhlhaas, 2014)) and theta range (5–15 Hz) (Jacobs & Kahana, 2009; Raghavachari et al., 2001) organize multi-item WM. Specifically, items are represented by the spatial pattern of neural activity within a gamma cycle; sequential list items are represented serially in consecutive gamma cycles, a process that repeats on each theta cycle (Fig. 1b). Such organization has been termed the theta-gamma code (Lisman & Jensen, 2013; Lisman & Buzsáki, 2008) and has been shown to be important in hippocampal processes that underlie encoding of episodic memory sequences (Heusser, Poeppel, Ezzyat, & Davachi, 2016) and the replay of spatial paths (Csicsvari, Jamieson, Wise, & Buzsáki, 2003; Gupta, van der Meer, Touretzky, & Redish, 2012; O’Keefe & Recce, 1993).

Although it is now clear that the theta-gamma code has a critical role in hippocampal function, there is only limited information about the involvement of this coding scheme in neocortex (Canolty et al., 2006; Rajji et al., 2016; Sauseng et al., 2009), the site of most WM processes (Milner, 2005). Computational models based on this coding scheme have been developed that fit the RT distributions in the Sternberg paradigm (Lisman & Jensen, 2013) and provide a plausible physiological account of the ordered serial representation of memory items in WM. However, testing this class of models has been difficult because of uncertainty about the cortical location of WM processes. In humans, localized oscillations can best be studied using intracranial electrocorticographic (ECoG) recording (Canolty et al., 2006), but this method has revealed different oscillation patterns at different sites during WM tasks, making it unclear which patterns subserve WM function (Raghavachari et al., 2001). Thus, it remains unclear whether CFC occurs in cortex during WM, as required by the LIJ model. Moreover, whether the theta-gamma coding scheme does in fact represent different WM items at different theta phases (i.e., serially in different gamma cycles) has not been previously possible to test.

We have studied the role of oscillations during WM by analyzing cortical sites that have involvement in WM, as demonstrated by stimulus-induced gamma-band power that depends on the identity of the letter presented. Such content-dependent sites, primarily in the occipital and temporal lobes, were identified in recent studies (Jacobs & Kahana, 2009; van Gerven et al., 2013), but their role in WM processes was not examined. Here we provide the first analysis of the properties of the oscillations at these sites during the encoding and maintenance phase of a WM task. The results reveal a complex regulation of both theta and gamma-band activity. Notably, the relative amplitudes of these oscillations are very different during encoding and maintenance. CFC is observed during maintenance. The presence of CFC during WM raised the possibility that this code might organize serial representations of WM items, and the existence of item-specific elevation of gamma power made it possible to test this possibility. The results show that the theta phase of gamma elevation during WM maintenance depends on the order at which the item was presented during encoding. These results support the hypothesis that multi-item WM involves serial activation of memories, as first hypothesized by Sternberg (S Sternberg, 1966) and physiologically implemented in the LIJ model (Fig. 1b).

## Results

Intracranial recordings were obtained while subjects performed the Sternberg WM task (a total of 1315 sites in 15 subjects). During the encoding phase of the task, three letters were presented sequentially, each visible for 700 ms with an inter-stimulus interval of 275–350 ms. This was followed by a 2-s maintenance phase. Finally, a probe letter was presented and subjects had to answer rapidly whether the probe was on the just presented list (Fig. 1a). As in previous work (Jacobs & Kahana, 2009; Rodriguez Merzagora et al., 2014; van Gerven et al., 2013), we first identified “*letter-selective*” sites as ones at which the increase in gamma-band power (70-100 Hz) depended on which letter was presented. As in Jacobs et al. (Jacobs & Kahana, 2009), letter-selective sites were rare (15 sites in 11 subjects; 4 subjects had 2 sites) and were located in the occipital and temporal lobes, mainly in the fusiform gyrus, a region implicated in letter analysis by other methods (Dehaene & Cohen, 2011) (Fig. S1). We used a measure of mutual information to quantify the distinguishability of individual letters (p<0.05, Bonferroni corrected; permutation test; see Methods under letter-selective sites). Per site, for the pair of letters with the largest mutual information, the letter with the greater gamma-band power was termed the “*tuned*” letter, whereas the other letter was termed the “*untuned*” letter.

### Time frequency analysis of encoding

To understand the role of oscillations in memory representations at these sites, we performed time-frequency analysis. Figure 2 shows the results by averaging over 11 subjects, considering only the site in each patient that was most or least-selective. We examined how low frequency oscillations were affected by letter presentation (encoding) at this class of sites. In contrast to the gamma power enhancement produced by letter presentation, theta power was transiently decreased after presentation of each letter at an offset of 0.5–1 s (Figs. 2a and 2b; one-tailed t-test, N=11; p<0.001 for average power at 7–12 Hz). Thus, theta and gamma power at letter-selective sites are differentially affected by letter presentation.

**Figure 2:**
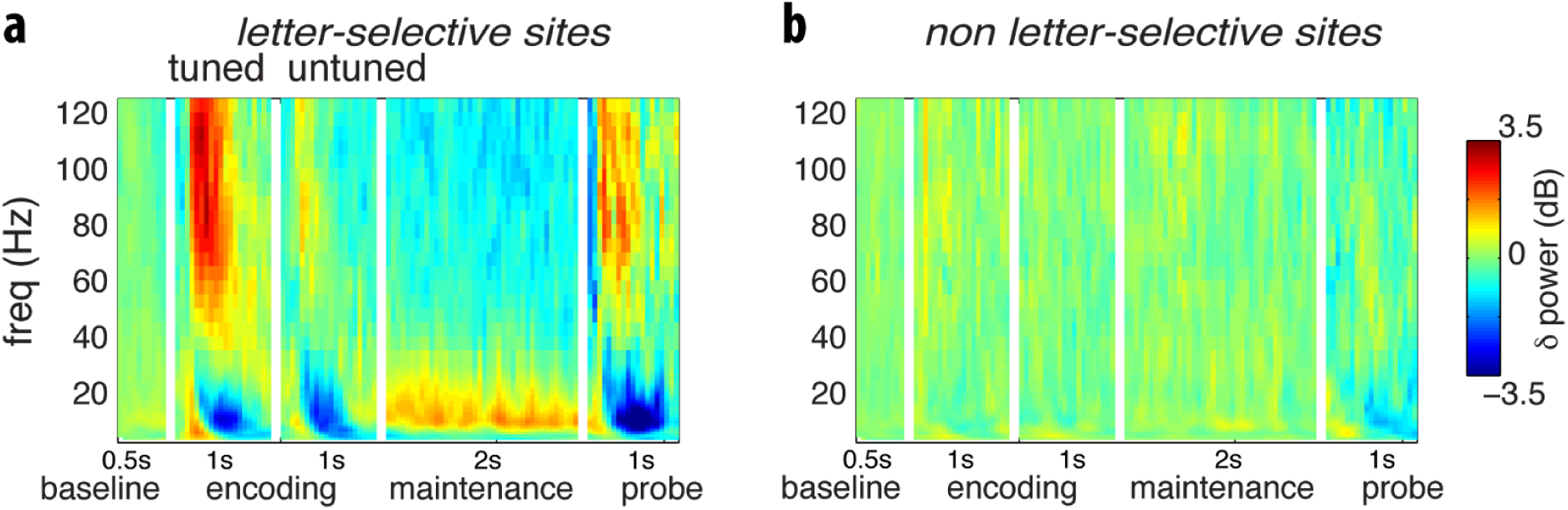
Time-frequency representations (TFRs) of power during encoding, maintenance, and probe. **(a)** Average TFRs of power at 11 letter-selective sites. The four subpanels are baseline responses to a turned letter, responses to an untuned letter, response during maintenance, and responses during the probe. During encoding, gamma is elevated by tuned letters; theta decreases by either the tuned or untuned letter. During maintenance, theta is elevated and gamma is very low. **(b)** Average TFRs of power at 11 non letter-selective sites shows little change during encoding or maintenance.

### Time frequency analysis of maintenance

We next turned to the analysis of the maintenance phase of the Sternberg task. As noted earlier, some previous work demonstrated continuous theta elevation during both encoding and maintenance (Raghavachari et al., 2001). As shown in Fig. 2a and 3c, this was not the case at letter-selective sites. Whereas theta power was greatly lowered during encoding, it became high during maintenance (Jacobs, Lega, Anderson, 2012)—indeed, significantly higher than baseline (N=15; two-tailed t-test p<0.01). Furthermore, whereas gamma-band power was higher than baseline during encoding, it was significantly lower than baseline during maintenance (N=15; two-tailed t-test p<0.01).

**Figure 3:**
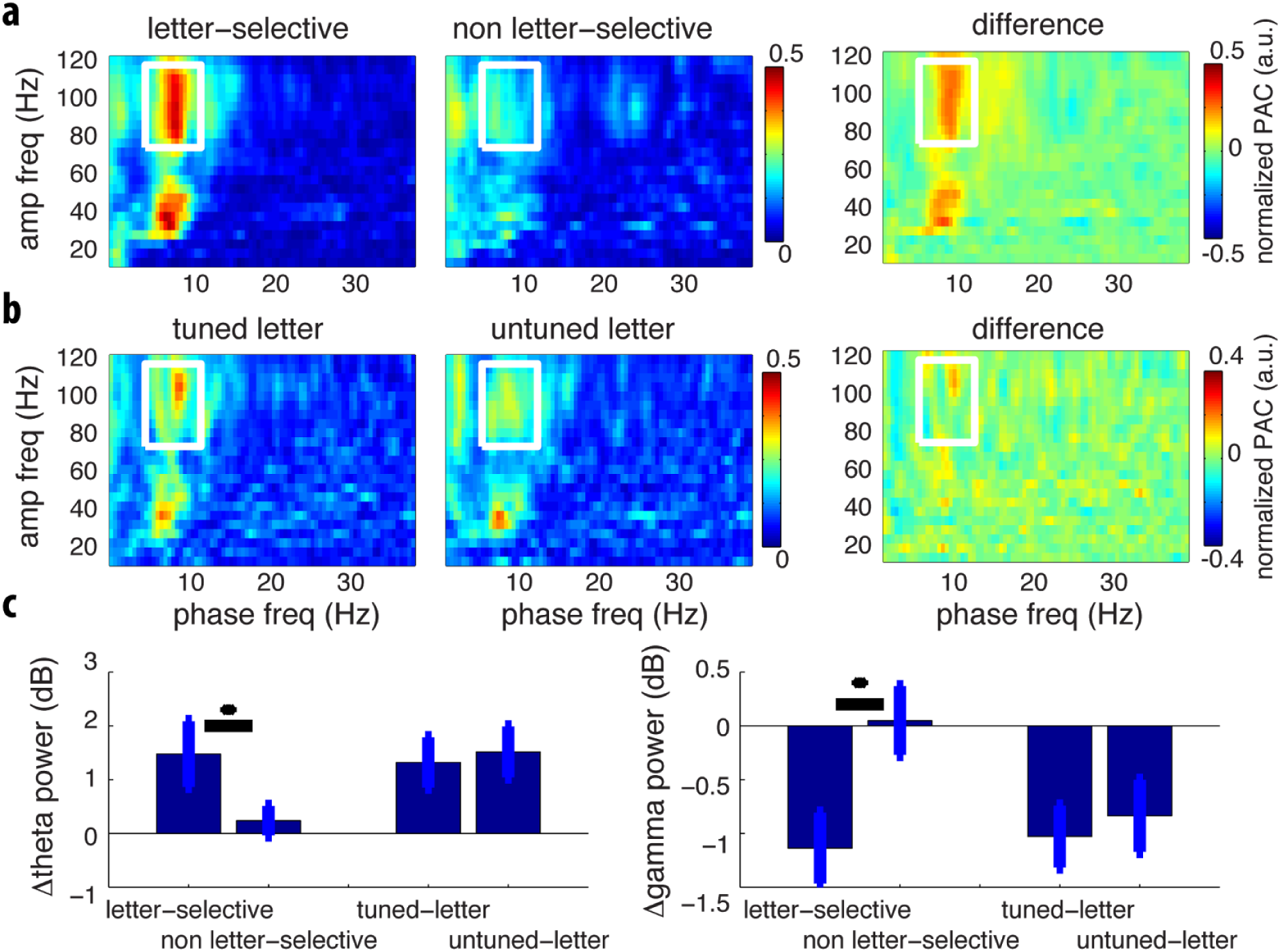
Properties of CFC during maintenance. **(a)** Average (11 sites) CFC is greater at letter-selective than at non letter-selective sites. In right panel, difference was calculated for each subject; the average difference is large, showing that CFC is much larger at letter-selective sites. **(b)** Average (15 sites) CFC is larger for tunes as compared to untuned letters. In panel at right, difference for each subject was calculated before averaging. **(c)** There is a significant difference (p<0.05) of theta and gamma power between letter-selective sites and non-letter-selective sites during the maintenance period. There is, however, no significant difference between theta and gamma power over letter-selective sites when comparing the tuned letter and untuned letter during the maintenance period.

Previous work on human cortical oscillations has shown that theta-gamma coupling, a form of cross-frequency coupling (CFC), can occur during cognitive tasks (Canolty et al., 2006; Rajji et al., 2016; Sauseng et al., 2009). It was of special interest to determine whether CFC occurs during the maintenance phase of this working memory task because this is a central assumption of the LIJ model. To quantify CFC, we used a measure of phase-amplitude coupling (PAC) that relates gamma-band power to the phase of low-frequency theta oscillations (Canolty & Knight, 2010; Jiang, Bahramisharif, van Gerven, & Jensen, 2015; see Methods under cross frequency coupling). The left panel of Fig. 3a shows the average normalized PAC over all letter-selective sites. During maintenance, there CFC between power in the 75–120-Hz range (fast gamma) and the phase of theta oscillations. This pattern was greater at letter-selective sites compared to non-letter-selective sites (N=11; one-tailed t-test p<0.05). It is noteworthy that the fast gamma range that showed CFC is the same range at which letter-selective gamma elevation was observed (Fig. 2a; see also Canolty et al., 2006).

If the same sites that represent items during encoding also participate in memory maintenance, then, oscillation properties at letter-selective sites should depend on whether a tuned item was actively held in memory after being viewed in the just presented list. Consistent with this expectation, CFC at letter-selective sites was greater when a tuned letter was on the just-viewed list (N=15; one-tailed t-test p<0.07; Fig. 3b; see Methods under test for normal distribution).

Our follow-up analyses confirmed that this pattern specifically reflected item-related changes in CFC. Theta- and gamma-band power (Fig. 3c right) were similar, irrespective of whether a tuned letter was just viewed (p’s>0.5), suggesting that the change in CFC that we observed for remembering a tuned letter during maintenance was unlikely to be a secondary result of power changes. It has been argued that non-sinusoidal waveforms can produce spurious CFC due to the power contributed from the higher harmonics (e.g., Aru et al., 2015). It is unlikely that this increased increased CFC for remembering a tuned letter (Fig. 3b) is explained by spurious coupling, as it was not associated with different overall theta or gamma power (Fig. 3c) (Jensen, Spaak, & Park, 2016).

### Testing Predictions of the Sternberg and LIJ models

In the Sternberg task, subjects make few errors, but exhibit a large variation in response time (RT). LIJ models that fit this large variation (Jensen & Lisman, 1998) hypothesize that RT variation is not due to processes during maintenance (problems during this phase would lead to errors) but, rather, due to processes during recall. Consistent with this, there was no significant relationship between CFC during maintenance and RT (p>0.3; N=15; Fig. 4). For a discussion of what determines RT, see Discussion.

**Figure 4:**
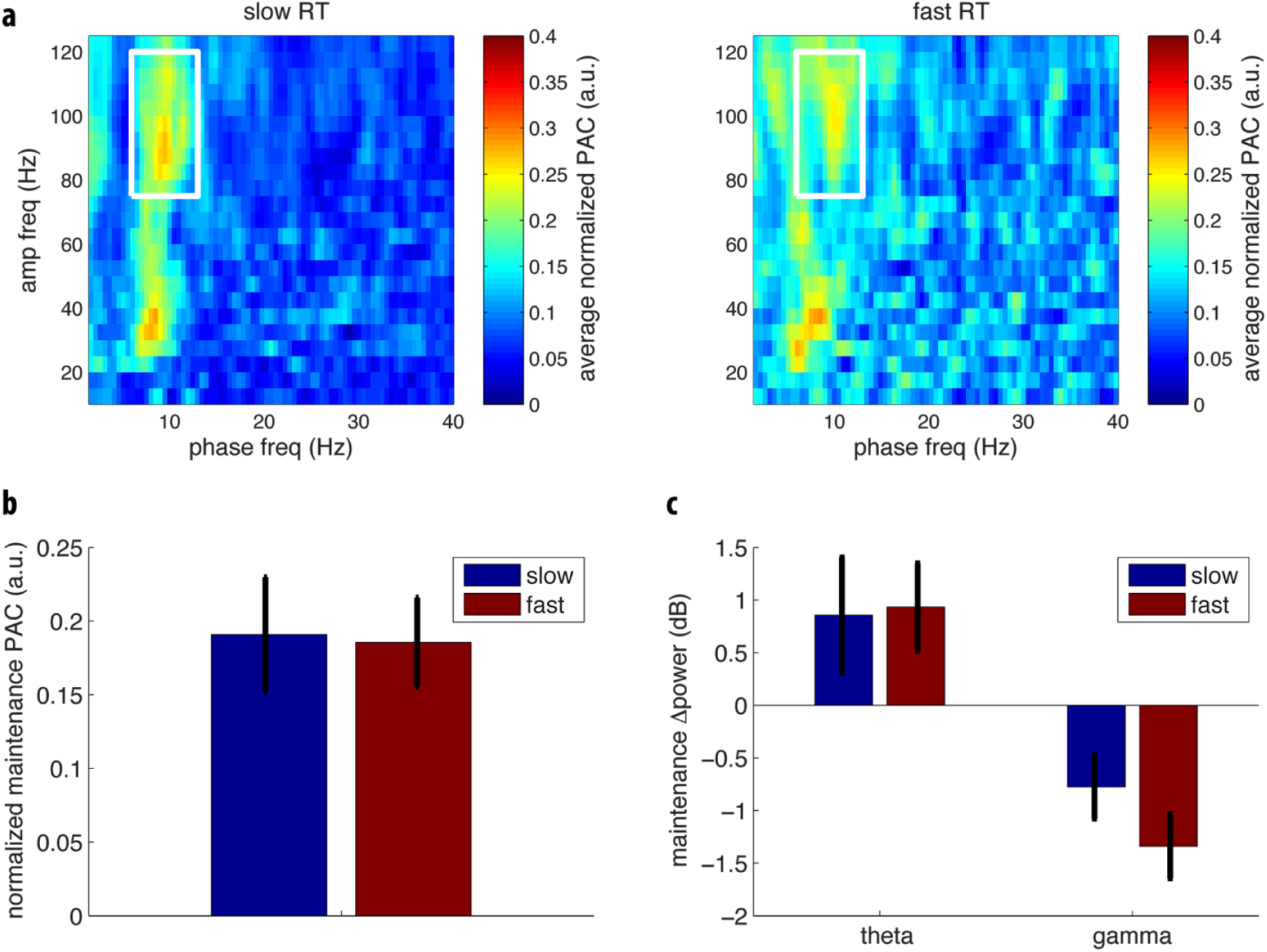
CFC during maintenance. **(a)** CFC in trials with slow or fast RT. **(b)** Average CFC (PAC) over the white box of panel a. **(c)** Average maintenance theta and gamma power. Error bars SEM.

A key prediction of the LIJ model is that different memory items are represented by neuronal activity at distinct theta phases (Fig. 1b). The broadband gamma signal that we measured reflects the mean spiking activity of individual neurons (Manning et al., 2009). Thus we predicted broadband gamma power at a letter-selective site would be elevated at the moment when a tuned letter is represented and, moreover, the theta phase of maximum power would vary with the serial list position of a tuned letter (Fig. 1b). We tested this prediction by measuring fast gamma-band power (75-120 Hz) for tuned letters presented at the three list positions (P1, P2, and P3). For each site, the position-specific deviation of fast broadband gamma-band power from the overall average was measured as a function of the local theta phase.

As shown in Fig. 5a, the preferred phase of the activation for tuned letters depended on their position in the list; broadband gamma-band power was significantly higher at an earlier phase for P1 than for P2 and at an earlier phase for P2 than for P3 (N=15; one-tailed t-test p<0.05; see Fig. 5a). A second way of analyzing this data, which confirmed the conclusion based on Fig. 5a, is shown in Fig. 5b. This figure shows average fast gamma power as a function of theta phase over sites, colored to indicate which item position (P1, P2, P3) had the maximum power. It can be seen that the color changes systematically. Using a permutation test, we found that this pattern, consisting of three discrete clusters, was unlikely to have occurred by chance (p<0.003; permutation test; see Methods under permutation test for serial ordering). A similar phase dependence was not observed for slow gamma (p>0.2; permutation test). Thus, item activation during memory maintenance, as measured by fast gamma-power, has a theta phase that depends on when the item was presented during encoding.

**Figure 5:**
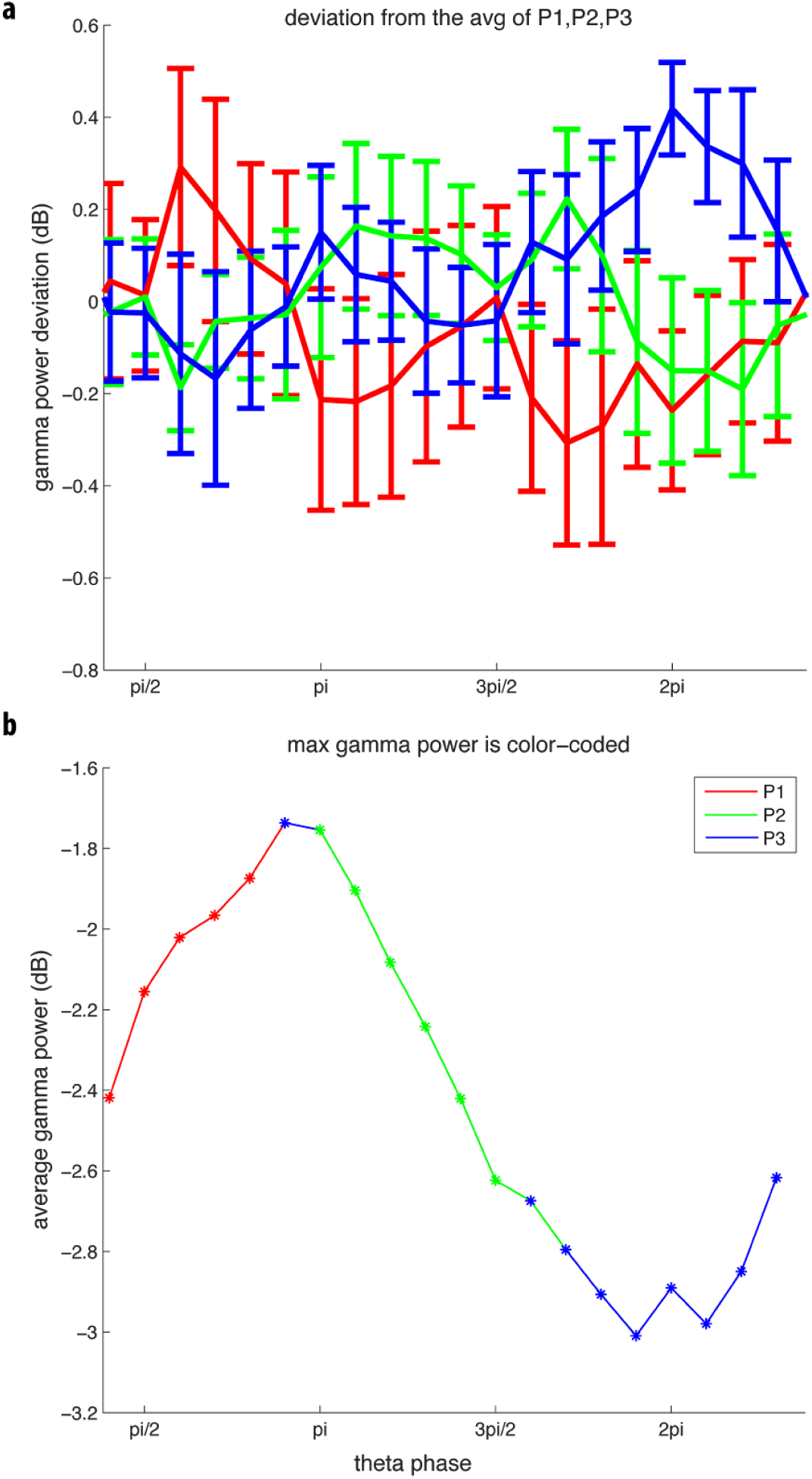
List position affects the theta phase that has maximal fast gamma power during maintenance. **(a)** The deviation of fast gamma-band power for each position from the average measured as a function of theta phase (Hilbert phase 7-13 Hz, see Methods under permutation test for serial ordering). Error bars represent SEM. **(b)** Average fast gamma power as a function of theta phase; the color indicates which item position (P1-P2-P3) yielded the maximum power at that phase bin.

## Discussion

Understanding human cortical WM processes has been complicated by the diversity of response patterns in different cortical sites, a diversity we confirm here. A key step in the effort to understanding WM was therefore the discovery (Jacobs & Kahana, 2009) of cortical sites with demonstrable mnemonic content, as indicated by letter-selective gamma elevation. Here we provide the first analysis of the multiple oscillatory processes at such sites. Our results show that the amplitudes of oscillations are strongly affected by the task. However, contrary to sites where a fixed oscillation pattern is gated on/off at the beginning/end of the task (Raghavachari et al., 2001), letter-selective sites showed very different oscillatory patterns during the encoding and maintenance phases of the task. Notably, high-frequency activity in the gamma range and low-frequency activity in the theta range were inversely affected during encoding/maintenance. Cross-frequency coupling (CFC) also occurred during maintenance. The ability to observe CFC during maintenance and the existence of letter-selective gamma elevation allowed us to test the concept of theta/gamma coding in WM. The results provide the strongest support to date for the serial organization of WM, as first proposed by Sternberg (S Sternberg, 1966), and for the proposal (LIJ model) that this organization depends on CFC mediated by a theta/gamma code (Fig. 1b).

### CFC during WM maintenance

Cross-frequency coupling between gamma oscillations and theta oscillations is easily observed in the local field potential of the cortex and hippocampus in rodents and humans (Axmacher et al., 2010; Bragin et al., 1995; Jacobs, Lega, & Anderson, 2012). According to the LIJ model, the function of these dual oscillations is that they form a general coding scheme (now called the theta-gamma code; reviewed in Lisman & Jensen, 2013; Lisman & Buzsáki, 2008) according to which each item is represented by the spatial pattern of firing in a network that occurs during a given gamma subcycle of a theta cycle (Fig.1b).

Evidence for such temporal coding organized by oscillations came from the discovery of theta-phase precession in hippocampal place cells (O’Keefe & Recce, 1993) and the evidence that gamma oscillations subdivide the theta cycle into discrete phases (Buzsáki, 2002; Gupta et al., 2012; Lisman & Buzsáki, 2008). Other work has shown that disruption or enhancement of theta-gamma oscillations can affect memory, pointing to the functional importance of this code (Shirvalkar, Rapp, & Shapiro, 2010). Intracranial recordings in humans have demonstrated that the hippocampus demonstrates CFC during working memory (Axmacher et al., 2010). Importantly, recent work shows that, during the encoding of item sequences in the human hippocampus, different list items are represented at different theta phases (Fuentemilla et al., 2010; Jafarpour et al., 2014; Heusser et al., 2016), providing support for the hypothesis that the hippocampus utilizes a theta-gamma code for episodic memory.

### Theta-gamma code for cortical WM processes

There has been slower progress is understanding the mechanisms generating oscillations in neocortex (Lee et al., 2005; Siegel, Warden, & Miller, 2009). In neocortex, theta oscillations are generated by input from the thalamus and basal forebrain (Jones, 2002), slow gamma activity is locally generated by interaction of pyramidal cells and interneurons (Börgers, Epstein, & Kopell, 2005), and fast gamma activity is largely generated by spiking activity (Belluscio et al., 2012; see also below). The existence of letter-selective sites in cortex has now made it possible to study temporal coding in more detail and to specifically test predictions of the LIJ model of WM (Fig. 1b). This model predicts that items are represented at different theta phase according to their order in the presented list. Thus, the specific prediction is that the elevation of fast gamma power produced by presenting a letter to which a site is optimally responsive (tuned) should occur at a theta phase that depends on the item’s order in the list. The results in Fig. 5 show this to be the case. This suggests that the fundamental solution to keeping multiple items active in WM is by a multiplexing mechanism in which different items activate at different phases of theta, i.e., serially.

A second prediction of the LIJ model is that the serial organization of different memories is organized in time by ongoing gamma oscillations. This leads to the question of which type of gamma oscillation has this role. As shown in Fig. 3a left panel, CFC of gamma and theta is present both in a broadband (fast gamma) range (70-120 Hz) and in a lower (slow gamma) range (30 to 50 Hz). There are now several lines of evidence that fast gamma arises from spiking activity, an activity that is temporally organized by slow gamma. Notably, broadband gamma correlates with spiking activity (Belitski et al., 2008; Buzsáki & Wang, 2012; Jacobs & Kahana, 2009; Lashgari et al., 2012; Liu & Newsome, 2006; Miller, Honey, Hermes, Rao, & Ojemann, 2014; Ray, Crone, Niebur, Franaszczuk, & Hsiao, 2008; Ray & Maunsell, 2011), spiking is phase locked to slow gamma (Jacobs, Kahana, Ekstrom, & Fried, 2007; Kraskov, Quiroga, Reddy, Fried, & Koch, 2007), and fast gamma shows CFC with slow gamma (Bahramisharif et al., 2015; Miller et al., 2010). Thus, the fact that fast gamma activity is item-specific (Fig. 2a) and has theta phase that depends on order (Fig. 5) suggests that it reflects the spiking of cells that represent particular list items. It is notable that the power of slow gamma does not display substantial letter selectivity (Fig. 2a). Thus the data suggest that slow gamma does not itself carry information but serves to temporally organize the firing of cells that carry information, as postulated in Fig. 1b.

### Relationship to behavioral results in the Sternberg task

The classical behavioral results on the Sternberg task were obtained in laboratory setting using well-trained subjects who were highly motivated to respond as quickly as possible. These results show that RT varies linearly with set size and has an average slope of 38 ms for simple list items (letters or numbers) (S Sternberg, 1966). The exact properties of the RT distributions are known. The patients from whom we obtained brain recordings were not trained or highly motivated and the experiments were done in a hospital setting. The RT in these experiments thus tended to be considerably slower than those obtained in the laboratory. However, it is nevertheless of interest to consider whether the underlying brain processes we have observed in patients might relate to the timing processes revealed in laboratory experiments. The experiments in healthy subjects have provided information not only on how the mean RT depends on set size (S), but also how the standard deviation and skewness depends on S. This data could be accounted for by either of two computational models (Jensen & Lisman, 1998). In the “adapting theta model,” theta frequency was assumed to change with load, a change recently observed in the hippocampus (Axmacher et al., 2010). The best fit of this model led to the prediction that the theta frequency for S=3 is 8.3 Hz. This frequency is in good agreement with the ∼9 Hz theta we have observed experimentally. In the adapting theta model, the Sternberg slope (38 msec) is 1.5-2 times the period of the underlying gamma oscillations. The model predicts a gamma frequency of 39-52 Hz. This is at least roughly consistent with the frequency range of slow gamma (30-50 Hz) that we observe in CFC plots (Fig. 3). A second model assumes that stimuli reset the phase of theta, a process also observed (Jutras, Fries, & Buffalo, 2013; Williams & Givens, 2003). This “phase reset” model gave an estimate of theta of 7.2 Hz and gamma of 47 Hz. Thus, these models, both of which are based solely on fits to RT distributions in the Sternberg task, make reasonable predictions about the critical frequencies observed in the ECoG data. This good correspondence suggests that the slow RTs observed in patients probably reflect motor/attention processes rather than memory retention processes. Now that it is clear how much can be learned from the study of letter-selective sites, it will be of great interest to study theta reset and and the effects of varying memory load. This should make it possible to develop more exact models of how oscillations organize WM.

A successful theory of WM should also be able to explain the remarkable behavioral findings of Cavanaugh (Cavanagh, 1972). He reported that the Sternberg slope increases with item complexity, and furthermore, that the number of items that could be remembered (span) decreases with item complexity. Remarkably, the product of span and slope remains constant. As argued in a recent review (Saul Sternberg, 2016), the LIJ framework could explain this constancy if it assumed that span is determined by the number of gamma cycles within a theta cycle and that gamma period increases with item complexity. Recent physiological findings are relevant here. If complex items are represented by less sparse activity, as seems reasonable, then the known dependence of gamma period on activity level (greater activity results in slower gamma; Atallah & Scanziani, 2009) could account for why gamma period increases with complexity. Experiments could now be conducted to test the relationship of gamma period to item complexity, potentially providing a mechanistic basis for Cavanaugh’s findings. More generally, if oscillations underlie working memory, notably the span, there should be a relationship between the two. Consequently, procedures that manipulate oscillations properties should affect span. Although such efforts are in their early stage, there are notable successes (Alekseichuk et al., 2016; Axmacher et al., 2010; Chaieb et al., 2015; Fukuda, Mance, & Vogel, 2015; Vandenbroucke et al., 2015; Vosskuhl et al., 2015).

### Other models of WM

How does the evidence for theta-gamma model relate to other models that offer a different view of WM mechanisms? In the classic studies of single unit activity during WM, firing persisted during the entire delay period (Funahashi et al., 1989). However, recent findings indicate that activity can be discontinuous rather than continuous (Baeg, Kim, Huh, & Mook-Jung, 2003; Noy et al., 2015; Stokes, 2015; Vugt, 2014; Watanabe et al., 2014). For example, sites in frontal cortex that have mnemonic activity have brief and sporadic bursts of firing and gamma oscillations (Lundqvist, Rose, & Herman, 2016). According to one theoretical framework (Lundqvist, Herman, & Lansner, 2011), these bursts may depend on item-specific changes in synaptic weights that occur during encoding. If information has been encoded in weights, it follows that persistent firing may not be necessary for the maintenance of stored information.

One possibility is that this evidence for discontinuous activity and the theta-gamma framework we have proposed are in fact compatible and reflect the existence of dual memory mechanisms (probably in different locations), as suggested by widely accepted psychological models that were based on memory tests in normal and amnesic patients (Atkinson & Shiffrin, 1968; Baddeley, 2000). According to these models, items are first stored in a cortical limited-capacity buffer. Output from this buffer then drives synaptic modification in the high-capacity hippocampal network. Depending on conditions, recall of lists held in WM may depend on either the cortical buffer or the hippocampus. The network based on the LIJ model has the properties required to implement the buffer. Importantly, it can rapidly store even novel information because information storage does not depend on synaptic modification, a process that can take seconds to develop (Gustafsson, Asztely, Hanse, & Wigstrom, 1989); rather, memories are reactivated on every theta cycle by activity-dependent intrinsic conductances. It seems possible that the sporadic gamma bursts described above might reflect the information stored in the hippocampal system. Indeed, experiments show that, once information is encoded in the hippocampal network, memories are sporadically reactivated during brief events termed sharp-wave ripple (SWR) (Watson & Buzsáki, 2015). Moreover, the SWR can drive activity in cortex (Headley, Kanta, & Pare, 2016; Remondes & Wilson, 2015). It will thus be of interest to test whether the sporadic gamma bursts observed in cortex during WM maintenance are the result of sporadic SWRs generated by the hippocampal system.

### Properties and function of low-frequency oscillations

Our results provide new information about the control of theta oscillations at content-specific cortical sites. As noted previously, various patterns of activity have been observed at sites not specifically linked to WM content (see also Sauseng et al., 2009). For instance, a previous study identified sites that undergo increases or decreases in theta power, changes that persist during both encoding and maintenance phases of the task (Raghavachari et al., 2001). Here we show a quite different pattern at letter-selective sites—theta power goes down during encoding but up (even above baseline) during maintenance. The suppression may be related to the spatially non-uniform suppression of theta power that occurs in visual and sensori-motor cortices when a subregion is involved in a task (Harvey et al., 2013; Pfurtscheller & Lopes da Silva, 1999).

We show here that during maintenance there are sites that show elevated theta power and that are nevertheless engaged in WM maintenance, as suggested by the fact that CFC is content-specific. Other data suggests that, when low-frequency power becomes high, it inhibits the processing of sensory inputs (Jensen & Mazaheri, 2010; Raghavachari et al., 2001) (this is generally termed alpha range, which we refer to here as theta). One way of reconciling these results is to suppose that theta (alpha) generators in cortex and thalamus (Bollimunta, Mo, Schroeder, & Ding, 2011) inhibit the flow of incoming sensory information during maintenance but do *not* inhibit and indeed protect (Bonnefond & Jensen, 2012; Payne, Guillory, & Sekuler, 2013) the information that is already held in WM buffers. On the other hand, the lowering of theta power during encoding may be permissive of sensory data entering working memory buffers.

## Conclusions

Although theta-gamma coupling appears in many brain regions (Igarashi, Isomura, Arai, Harukuni, & Fukai, 2013), the generality of the theta-gamma code remains unclear (Lisman & Jensen, 2013). Early work established that this code organizes spatial information in the hippocampus (Csicsvari et al., 2003; Heusser et al., 2016; Igarashi et al., 2013; O’Keefe & Recce, 1993), and recent work on the hippocampus has shown that it also organizes non-spatial information (Heusser et al., 2016). Here, by focusing on sites demonstrably engaged in WM, we have shown that the theta-gamma coding occurs in cortex; specifically, the theta phase of item representation depends on item order during presentation. Furthermore, the estimates of theta and gamma frequencies that we obtained by measurement are in reasonable agreement with a model of RT distributions based on purely behavioral results. Thus, these results provide a coherent view of how the theta-gamma code could mediate WM. Taken together, these findings point to the possibility that the theta-gamma code is a general coding scheme appropriate for representing ordered information in both cortex and hippocampus.

## Methods

### Recording

Data of 15 epileptic patients (6 females, age: 36 +/- 9) implanted with electrocorticography (ECoG) grids were included in this study. All subjects performed a Sternberg-type working memory task (S Sternberg, 1966) in which three letters were presented during encoding, and after 2 seconds of retention, a probe letter was presented either from the memory list or not. The research protocol was approved by Institutional Review Boards at the Hospital at the University of Pennsylvania (Philadelphia, PA) and the Thomas Jefferson University Hospital (Philadelphia, PA). Informed consent was obtained from each patient or their legal guardians. The details of the experiment and how data was collected have been previously reported in another paper (Rodriguez Merzagora et al., 2014) where data from 15 patients were selected for inclusion based on sensor coverage and trial counts. The included data was selected at the beginning of this study prior to data analysis. Our reasons for excluding data from the earlier study (Rodriguez Merzagora et al., 2014) included the following: 1. Some data came from an unsuitable variant of the task with a short maintenance period. 2. The clinical recordings for some subjects contained transient interruptions due to file changes. 3. Some subjects had been recorded at a low sampling rate that precluded examining CFC. No data was collected during a seizure, and none of the analyzed sites with letter selectivity had been designated as sites where there had been seizure activity.

### Artifact Rejection

We distinguished sites that were involved in seizure initiation, based on information provided by clinical teams. We then used kurtosis to identify EEG segments with epileptiform discharges (Guggisberg et al., 2004; Robinson et al., 2004, Sederberg et al., 2007; van Vugt et al., 2010). We computed the kurtosis for each individual EEG segment and identified any segments with kurtosis above 5 as being potentially epileptogenic. Visual inspection confirmed that this approach worked correctly, because these periods, although rare, contained unusual EEG such as epileptic spikes or sharp waves. For more detail on this method see van Vugt et al. (2010).

### Preprocessing

All recordings were resampled to 250 Hz. Further, they were re-referenced to the common average over all electrodes. Line noise was removed by using discrete Fourier transform filtering centered at 60 and 120 Hz.

### Statistical Assessments

Two-tailed t-test was used for the majority of statistical assessments. In case of using one-tailed t-test or permutation test, it is clearly stated next to the p-value. The degree of freedom (df) was always set to N-1 where N was the number of samples.

### Time Frequency Analysis

Time-frequency analysis of phase and power was performed using a sliding window approach. To estimate the power, we used time windows of three cycles of low-frequency oscillations (from 1 to 30 Hz in step of 1 Hz) and a fixed length of 100 ms for high-frequency oscillations (from 30 to 125 Hz in step of 5 Hz) centered at -1 to 7 seconds relative to the first letter presentation with the step size of 50 ms. The high-frequency power estimate may have included both gamma oscillations and other high-frequency neural activity (Manning & Jacobs, 2009). Prior to the FFT, the windows were multiplied with a Hanning window of the same length.

For each trial, baseline power was calculated as the average power over the bins from -0.8 to -0.35 s relative to the onset of the first letter presentation. For the encoding period, we used 0.2 to 0.5 s relative to the onset of each letter, and for the maintenance period, we used -1.85 to -0.3 s relative to the probe onset. The group statistics were performed using t-test.

### Letter-selective sites

We used mutual information as the measure of dependence between gamma power and the viewed letter. This procedure identified, across all possible letter pairs X and Y, sites that exhibited significant mutual information distinguishing when the patient was viewing one letter (X) compared to another (Y). We computed this measure as: I(X;Y)= H(X)-H(X|Y) where H is the entropy function. In our paper, X is the baseline-corrected gamma power (70–100 Hz) and Y is a binary vector where 0 stands for the first letter in the pair and 1 stands for the second letter. In our case of having 16 letters, there were 16x15/2=120 letter pairs that tested for each electrode. For the statistical assessment, a null distribution was made by randomly permuting class labels and recalculating the mutual information. We Bonferroni corrected the p-value for the number of electrodes and 120 letter pairs.

For each electrode, the pair of letters that resulted in the highest mutual information was selected, and the one with the higher gamma power was termed the *tuned* letter and the other one was termed the *untuned* letter. Therefore, when a tuned letter was presented to the patient, the relevant letter-selective site exhibited an increase in gamma power during encoding. This pattern was significant (p<0.05, Bonferroni corrected; permutation test) in 15 independent sites in 11 subjects out of 15. We consider each electrode independently because multiple sites in any given patient were more than 1 cm apart and the highest mutual information across these sites was always obtained for different pairs of letters.

To examine letter tuning of content-specific sites during maintenance, for the tuned letters, we only considered trials in which at least one of the maintained items was the tuned letter and where the untuned letter was not viewed. Inversely, for examining the maintenance of untuned letters at these sites, we considered trials in which at least one of the remembered items was the untuned letter and none of them was the tuned letter.

### Cross-frequency coupling (CFC)

We used ‘phase-to-amplitude coupling’ (PAC) to investigate CFC. PAC was calculated as the coherence between low-frequency oscillations and the amplitude of high-frequency activity (Jiang et al., 2015). For each trial, the power-envelope of the high-frequency activity was estimated from the FFT of a frequency dependent sliding time window (six cycles long; i.e. dT = 6/f) multiplied by Hanning taper. This was done from 10 to 125 Hz in steps of 5 Hz. The CFC was calculated from the coherence between this envelop and the signal. The 2 seconds of the retention interval was divided into three windows of 1s (centred at 0.5 s, 1.0 and 1.5 s in the retention interval; i.e. 50% overlap). Using the same methods as described in (Jiang et al., 2015), the coherence was calculated over these time-windows and trials. Note that the trials were not concatenated, but we calculated the PAC for individual trials and then averaged.

When comparing CFC values in the maintenance period, for each subject, the PAC values were averaged (phase frequency of 7–13 Hz; amplitude frequency 75–120 Hz). These frequency ranges are consistent with the dominant frequency values in Figure 2. To compare CFC over sites, per subject, the average PAC values over all trials were divided from the letter-selective site by the non-selective site (11 sites). Furthermore, over letter-selective sites, the average PAC values over trials with tuned letters were divided by the average over trials with untuned letters (15 sites). Using paired t-tests, we tested whether the values obtained are greater than 1 over subjects. Because the statistics were done on ratios, the normalization, which was used for visualization in Fig. 3, did not affect the statistical tests. In several situations we reported a single PAC value per electrode rather than a spectrum. Here we first computed the mean level of PAC for each subject in the frequency range of interest. Having identified these individual PAC values, we then performed statistical analyses across this distribution. This frequency window was predefined consistent with the literature and Figure 2 (7–13 Hz by 75–120 Hz), thus avoiding concerns on multiple comparison. Note that all statistical tests were performed on the raw PAC measures in this window using paired tests, with the normalization being performed for visualization purposes only.

### Test for normal distribution

To test whether the distribution of CFC ratios follows a normal distribution, we first performed Lillifors’ normality test. The results were consistent with the distribution being normal (p<0.05). We then generated a set of normally distributed values using the same mean and standard deviation of our ratios and used Kolmogrov-Smirnoff test to evaluate whether the two distributions are equal. After 1000 iterations, the null hypothesis that the ratios are *not* normally distributed was rejected (p<0.01). On the basis of these analyses we concluded the use of t-test was valid.

### CFC normalization for visualization

First, we computed the complete PAC coupling time-frequency analysis for each electrode/subject. Then we normalized each individual PAC spectrum by dividing by the maximum observed PAC value, thus ensuring that each spectrum was the same scale by having the same maximum value. These normalized PAC spectra were then averaged over sites.

### Permutation test for serial ordering

Theta phase was calculated as the angle of the Hilbert transform of the data filtered at 7-13 Hz. Fast gamma power (75-120 Hz) during the maintenance period was estimated by applying the Hilbert transform to a Hanning-tapered sliding window of 40 ms. For the maintenance period of each trial, the 2π radians of theta phase were divided into 10 equal phase bins. Then for each of these 10 phase bins, the average fast gamma power was calculated. We performed this analysis for the maintenance period when the tuned letter was presented in the first (P1), second (P2), and third (P3) positions. Then for each phase bin per letter-selective site, we identified the list position with the highest fast gamma power. This analysis resulted in a sequence of 10 numbers for each site where each number can be 1, 2, or 3, referring to whether P1, P2, or P3 had the highest fast gamma power, respectively. Starting from theta phase zero, for the average normalized fast gamma power, we got the following sequence: “3311122233” (note that it is a circular sequence; see Fig. 5b).

For the statistical assessment, we determined the proportion of times when shuffled data provided this type of sequential pattern in the representation of the remembered item across theta phase. First we made 30 reference distributions where we divided the 10 bins into 3 equal divisions (base templates “1112223333”, “1112222333”, and “1111222333”, along with each of their 10 circular rotations). We defined a measure for the ‘distance’ between the observed pattern and the reference distribution as follows: we identified the template most similar to observed pattern. The ‘distance’ was then the number of bins being different (e.g., the distance between ‘”1112222333” and “1132221332” is 3). We formed a null distribution by randomly switching all labels of each site across trials and then performed the average distribution. We performed the randomization 10,000 times and counted how many times the distance was less than or equal to the distance of the sequence observed.

## Supplementary figure

**Figure S1:**
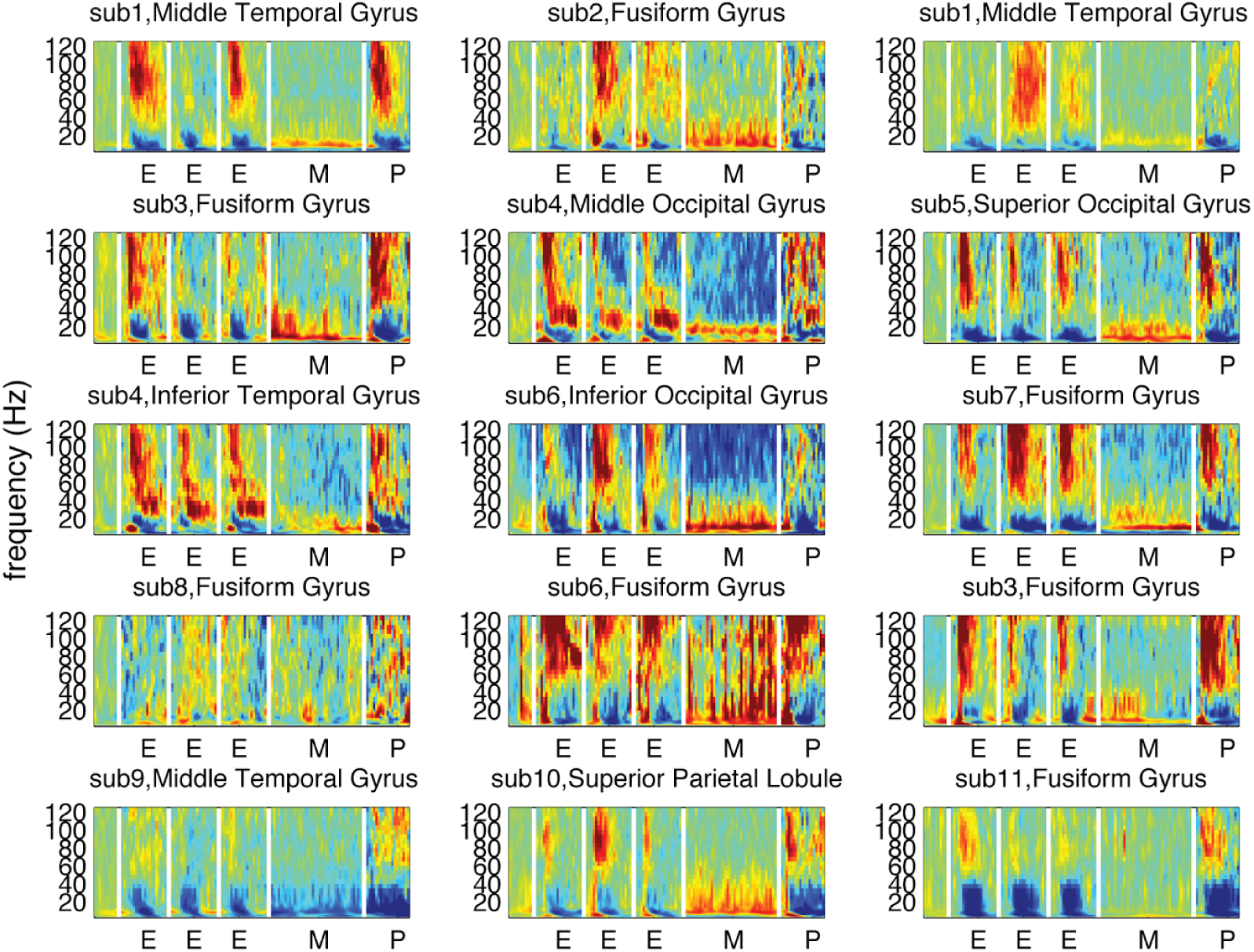
Demonstration that data from Fig. 2a is reproducible over the 15 letter-selective sites of 11 subjects. Time-frequency representation of power of the most selective and the least selective sites (highest and lowest mutual information) with their corresponding letters over 11 subjects. Average baseline power is subtracted from all time points. Subject number and location of electrodes are shown above each plot. E, M, and P refer to encoding, maintenance, and probe, respectively. The segments between white lines should not be interpreted as letter order. During different segments, letters with different tuning are presented.

## Acknowledgements

We acknowledge Nikolai Axmacher, Andrew Watrous, and Mike X. Cohen for giving constructive feedback on our paper. We thank Mikael Lundqvist and Mark Stokes for discussions about non-persistent activity during WM maintenance. We thank Gyuri Buzsaki, Paul Verschure, and Bart Gips for discussions about oscillations. O.J. acknowledges support from VICI grant (453-09-002) from The Netherlands Organization for Scientific Research (NWO) and a James S. McDonnell Foundation Understanding Human Cognition Collaborative Award 220020448. J.J. acknowledges support from NIH grants MH104606 and MH061975. J.L acknowledges support by NIH grants R01MH102841, U01NS090583, and R56NS096710.

